# Sex Differences in Sleep Phenotypes in the BACHD Mouse Model of Huntington’s Disease

**DOI:** 10.1101/2023.04.28.538324

**Authors:** Emily Chiem, Kevin Zhao, Gemma Stark, Cristina A. Ghiani, Christopher S. Colwell, Ketema N. Paul

## Abstract

Sleep and circadian rhythm disturbances are common features of Huntington’s disease (HD). HD is an autosomal dominant neurodegenerative disorder that affects men and women in equal numbers, but some epidemiological studies as well as preclinical work indicate there may be sex differences in disease progression. Since sex differences in HD could provide important insights to understand cellular and molecular mechanism(s), we used the bacterial artificial chromosome transgenic mouse model of HD (BACHD) to examine whether sex differences in sleep/wake cycles are detectable in an animal model of the disease. Electroencephalography/electromyography (EEG/EMG) was used to measure sleep/wake states and polysomnographic patterns in young adult (12 week-old) male and female wild-type and BACHD mice. Our findings show that male, but not female, BACHD mice exhibited increased variation in phases of the rhythms as compared to age and sex matched wild-types. For both Rapid-eye movement (REM) and Non-rapid eye movement (NREM) sleep, genotypic and sex differences were detected. In particular, the BACHD males spent less time in NREM and exhibited a more fragmented sleep than the other groups. Both male and female BACHD mice exhibited significant changes in delta but not in gamma power compared to wild-type mice. Finally, in response to a 6-hrs sleep deprivation, both genotypes and sexes displayed predicted homeostatic responses to sleep loss. These findings suggest that females are relatively protected early in disease progression in this HD model.

**Significance Statement:** Sleep and circadian rhythm disturbances are common features of Huntington’s disease (HD). Using an animal model of HD and EEG measures of sleep, we found that the males were more vulnerable to sleep/wake architecture alterations and sleep fragmentation. Homeostatic recovery from sleep loss did not appear to be altered at this stage of the disease. These findings raise the possibility that sex-specific factors play a role in the HD symptom progression and that hormone-based treatments may have therapeutic utility.

## Introduction

Sleep disorders are a common feature of many neurodegenerative diseases, such as Huntington’s disease (HD). Disruptions in the sleep/wake cycles of HD patients manifest early in disease progression, and are characterized by increased wakefulness during the night, daytime sleepiness, and delayed sleep onset (Morton, 2013; Fifel and Videnovic, 2020; Colwell, 2021). There is evidence that the suprachiasmatic nucleus (SCN), the central circadian clock, shows signs of extensive degeneration in HD patients (van Wamelen et al., 2014). Mouse models of HD also exhibit disruptions in the circadian rest/activity cycles similar to those observed in affected individuals. For example, the R6/2 mouse model of HD displays disruptions in the diurnal sleep/wake cycle, and increased sleep fragmentation (Morton et al., 2005; Fisher et al., 2013; Kantor et al., 2013). Similar changes in sleep behavior were seen in the Q175 mouse model, which shows increased wakefulness and decreased NREM sleep during the light phase (Loh et al., 2013; Fisher et al., 2016). The bacterial artificial chromosome transgenic mouse model of HD (BACHD), which offers the advantage of carrying the human mutation (Gray *et al*., 2008), also exhibits disruptions in the sleep/wake cycle (Kudo et al., 2011; Kuljis et al., 2016).

Sex differences in the development and manifestation of neurodegenerative diseases can offer important insights into disease pathogenesis. However, the literature on sex differences in HD patients and mouse models remains complex and limited. While some studies report sex differences in the age of onset, progression, and severity of the disease (Foround et al., 1999; Zielonka et al., 2013), others show no effect of sex (Wexler et al., 2004; van Dujin et al., 2014; Dale et al., 2016). There have been more consistent findings of sex differences in the preclinical models including the CAG 140 (Dorner *et al*., 2007), the Q175 (Padovan-Neto et al., 2019), and the BACHD mouse models (Kuljis et al., 2016). In HD patients, the lack of definitive evidence of sex differences may be due to the fact that such effects on these various components are influenced by other genetic or hormonal factors (Kehoe *et al*., 1999; Bode *et al*., 2008). Importantly, there has not been extensive investigation of how sleep/wake cycles differ by sex during the early stages of HD progression.

Sex differences in sleep/wake architecture and sleep homeostasis are well-characterized in the wild-type (WT_ mice (Paul et al., 2006; Paul et al., 2009). Hence, the present study was focused on determining whether sex differences can be detected in sleep architecture and homeostasis in young adult (12 week-old) wild-type and BACHD mice as measured using EEG.

## MATERIALS AND METHODS

### Animals

All the experimental protocols used to collect the data used for the present report were approved by the UCLA Animal Research Committee and followed the guidelines and recommendations for animal use and welfare set by the UCLA Division of Laboratory Animal Medicine and National Institutes of Health. The BACHD mouse model of HD used in this study contains a human mutant Htt gene encoding 97 glutamine repeats. BACHD females backcrossed on a C57BL/6J background were bred in house with C57BL/6J (wildtype, WT) males from Jackson Laboratory to obtain male and female offspring, either WT or heterozygous for the BACHD transgene. The WT littermates were used as controls. Both 3-month-old male (WT, n=14; BACHD, n=16) and female (WT, n=12; BACHD, n=13) mice were used in order to further our knowledge on the sex difference. Mice that had severe artifacts in their recordings were partially excluded from the analysis. Animals were group housed (4 per cage), and entrained to a 12:12 light-dark (LD) cycle, in sound-proof, humidity-controlled chambers until experimentation began.

### Surgery

All animals were surgically implanted at 12 weeks of age with EEG and EMG electrodes for polysomnographic recordings. A prefabricated head mount (Pinnacle Technologies, Lawrence, KS) was used to position three stainless-steel epidural screw electrodes. The first electrode (frontal-located over the front cortex) was placed 1.5 mm anterior to bregma and 1.5 mm lateral to the central suture. The second two electrodes (interparietal-located over the visual cortex and common reference) were placed 2.5 mm posterior to bregma and 1.5 mm on either side of the central suture. The resulting two leads (frontal-interparietal and interparietal-interparietal) were referenced contralaterally. A fourth screw served as ground. Silver epoxy was used to aid electrical continuity between the screw electrode and head mount. Stainless-steel teflon-coated wires were inserted bilaterally into the nuchal muscle to record EMG activity. The head mount was secured to the skull with dental acrylic.

### EEG/EMG recording

One week after surgery, mice were moved to sound-proof sleep-recording chambers and connected to a lightweight tether attached to a low-resistance commutator mounted over the cage (Pinnacle Technologies). Mice were allowed free range of movement throughout the cage while being tethered. Mice were given one week to acclimate to the tether and recording chambers. EEG and EMG recordings began at zeitgeber time (ZT) 0 (light onset) and continued for 24-hr. Data acquisition was performed on a personal computer running Sirenia Acquisition software (Pinnacle Technologies), a software system specific to rodent polysomnographic recordings. EEG signals were low-pass filtered with a 40-Hz cutoff and collected continuously at a sampling rate of 400 Hz. After data collection, waveforms were scored by the same trained operator as wake (low-voltage, high-frequency EEG; high-amplitude EMG), NREM sleep (high voltage, mixed frequency EEG; low-amplitude EMG), or REM sleep (low-voltage EEG with a predominance of theta activity (4-8Hz); low amplitude EMG). EEG epochs containing artifacts due to scratching, moving, eating, or drinking were excluded from analysis. Recordings were scored in 10-sec epochs. The operator was masked to the sex and genotype of the mice.

### Signal Analysis

Spectral analysis was performed on the frontal-interparietal lead. Power spectral analysis was performed by applying a fast Fourier transform (FFT) to raw EEG waveforms. Only epochs classified as NREM sleep were included in this analysis. Delta power was measured as spectral power in the 0.5-4 Hz frequency range and expressed as a percentage of total spectral power in the EEG signal (0.5-40 Hz). Power in the 0.5-4 Hz range (delta) was then averaged for all NREM epochs in a 24h period. Gamma power was measured as spectral power in the 20-40 Hz frequency range and expressed as a percentage of total spectral power in the EEG signal (0.5-40 Hz). Any changes in spectral power seen in the mid-active phase (ZT16-ZT22), were defined as changes in the siesta period. Sleep fragmentation was measured by the number of NREM bouts, duration of NREM bouts, and number of stage shifts.

### Total sleep deprivation

Immediately following a 24-hr baseline recording, mice underwent 6-hr of total sleep deprivation (SD) using a gentle-handling protocol, which includes cage tapping, introduction of novel objects, and delicate touching when mice displayed signs of sleep onset. SD began at the onset of the light phase in a 12hr:12hr LD cycle. Recordings continued for 18h of recovery sleep following the period of forced wakefulness.

### Statistical Analysis

Data were analyzed using SigmaPlot (version 14.5; SYSTAT Software, San Jose, CA). All values are reported as group mean ± standard error of the mean (SEM). The waveforms under baseline or after SD were analyzed by three-way analysis of variance (ANOVA) with genotype, sex and time as factors. The total time spent in each state (wake, REM, NREM), the amplitude, or phase of the rhythms, were analyzed by two-way ANOVA with genotype (WT, BACHD) and sex (male, female) as factors. We followed up with Bonferroni *post hoc* analyses when appropriate. Between-group differences were determined significant if p < 0.05.

## Results

### Wake rhythms

Prior work has found evidence for sex differences in overall activity rhythms in the BACHD mouse model of HD (Kuljis et al., 2016). In this study, we sought to use EEG measurements to determine whether these sex differences could be seen in daily rhythms of sleep/wake architecture early in the disease progression. Therefore, we tested the hypothesis that there are sex differences in the temporal patterns of wake in BACHD and WT mice at 12 week of age. The total amount of wake did not vary with genotype or sex (**Table 1**). Next, we examined the rhythm in wake using a three-way ANOVA with genotype, sex, and time (day, night) as factors. Both genotype and time had significant effects on the rhythms but not sex (**Table 2**). Examination of the waveform of this cohort using 2-hr bins revealed that the wake levels in BACHD males, but not females, were altered at several time points (**Fig. 1A, B; Table 3**). We quantified the amplitude of day/night difference as a ratio of minutes of wake in the night over the minutes measured during the day. While most mice had a similar amplitude, a subset of BACHD males exhibited a dramatically reduced amount of wake during the night (**Fig. 1C**). There was a significant interaction between genotype and sex in the rhythm amplitude (**Table 4**). Finally, BACHD mice exhibited a significant difference in the phase of peak wakefulness (**Fig. 1D, Table 4**). Strikingly, the male BACHD exhibited a great deal of variability in phase (**Table 5**) compared to the other 3 groups.

**Table 1:**
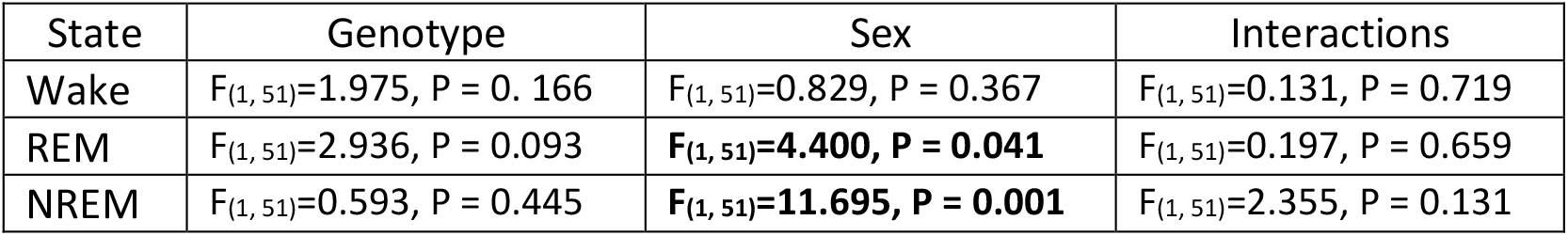
Analysis of total time spent in each state (wake, REM, NREM) using two-way ANOVA with genotype (WT, BACHD) and sex (male, female) as factors. In this and the other tables, significant values are indicated by bold.

| State | Genotype | Sex | Interactions |
| --- | --- | --- | --- |
| Wake | $F_{(1, 51)}=1.975$, $P = 0.166$ | $F_{(1, 51)}=0.829$, $P = 0.367$ | $F_{(1, 51)}=0.131$, $P = 0.719$ |
| REM | $F_{(1, 51)}=2.936$, $P = 0.093$ | **$F_{(1, 51)}=4.400$, $P = 0.041$** | $F_{(1, 51)}=0.197$, $P = 0.659$ |
| NREM | $F_{(1, 51)}=0.593$, $P = 0.445$ | **$F_{(1, 51)}=11.695$, $P = 0.001$** | $F_{(1, 51)}=2.355$, $P = 0.131$ |

**Table 2:** Analysis of waveforms using three way ANOVA with genotype (WT, BACHD), sex (male, female), and time (day, night) as factors. Interactions reported for genotype x sex x time.

| State | Genotype | Sex | Time (day vs. night) | Interactions |
| --- | --- | --- | --- | --- |
| Wake | $F_{(1, 107)}= 3.846$, $P = 0.053$ | $F_{(1, 107)}=0.433$, $P = 0.512$ | **$F_{(1, 107)}=251.196$, $P < 0.001$** | $F_{(1, 107)}=3.730$, $P = 0.056$ |
| REM | $F_{(1, 109)}=3.220$, $P = 0.076$ | **$F_{(1, 109)}=4.827$, $P = 0.030$** | **$F_{(1, 109)}=398.409$, $P < 0.001$** | $F_{(1, 109)}=0.202$, $P = 0.654$ |
| NREM | $F_{(1, 109)}=1.920$, $P = 0.169$ | **$F_{(1, 109)}=164.514$, $P < 0.001$** | **$F_{(1, 109)}=16.760$, $P < 0.001$** | $F_{(1, 109)}=0.283$, $P = 0.596$ |

**Table 3:** Analysis of waveforms using three way ANOVA with genotype (WT, BACHD), sex (male, female), and time (2 hr bins) as factors. In this table, significant values are indicated by bold. Interactions reported for genotype x sex x time.

| State | Genotype | Sex | Time (2 hr bins) | Interactions |
| --- | --- | --- | --- | --- |
| Wake | **$F_{(1, 599)}=13.921$, $P < 0.001$** | $F_{(1, 599)}=0.777$, $P = 0.379$ | **$F_{(11, 599)}=115.770$, $P < 0.001$** | $F_{(11, 599)}=1.788$, $P = 0.053$ |
| REM | **$F_{(1, 659)}=9.176$, $P = 0.003$** | **$F_{(1, 659)}=13.745$, $P < 0.001$** | **$F_{(11, 659)}=108.426$, $P < 0.001$** | $F_{(11, 659)}=0.460$, $P = 0.927$ |
| NREM | **$F_{(1, 635)}=4.380$, $P = 0.037$** | **$F_{(1, 635)}=21.725$, $P < 0.001$** | **$F_{(11, 635)}=89.398$, $P < 0.001$** | $F_{(11, 635)}=1.419$, $P = 0.160$ |

**Table 4:** Analysis of amplitude (ratio) and peak phase of each state (wake, REM, NREM) using two-way ANOVA with genotype (WT, BACHD) and sex (male, female) as factors.

| State |  | Genotype | Sex | Interactions |
| --- | --- | --- | --- | --- |
| Wake | Amplitude | $F_{(1, 49)}=3.521$, $P = 0.067$ | $F_{(1, 49)}=0.001$, $P = 0.970$ | **$F_{(1, 49)}=7.858$, $P = 0.007$** |
| | Phase | **$F_{(1, 51)}=0.002$, $P = 0.961$** | $F_{(1, 51)}=0.836$, $P = 0.365$ | $F_{(1, 51)}=1.323$, $P = 0.255$ |
| REM | Amplitude | $F_{(1, 49)}=0.610$, $P = 0.439$ | $F_{(1, 49)}=0.021$, $P = 0.885$ | $F_{(1, 49)}=0.002$, $P = 0.965$ |
| | Phase | $F_{(1, 51)}=1.851$, $P = 0.180$ | $F_{(1, 51)}=0.180$, $P = 0.673$ | **$F_{(1, 51)}=8.662$, $P = 0.005$** |
| NREM | Amplitude | $F_{(1, 49)}=0.120$, $P = 0.730$ | **$F_{(1, 49)}=17.727$, $P < 0.001$** | $F_{(1, 49)}=0.033$, $P = 0.856$ |
| | Phase | $F_{(1, 51)}=1.851$, $P = 0.180$ | $F_{(1, 51)}=0.180$, $P = 0.673$ | **$F_{(1, 51)}=8.662$, $P = 0.005$** |

**Table 5:** The confidence intervals in the measured of peak phase of the EEG rhythms by genotype and sex. The BACHD males exhibited strikingly more variable measures for each state.

| State | Genotype | Sex | Confidence interval |
| --- | --- | --- | --- |
| Wake | WT | Male | 0.966 |
|  |  | Females | 1.117 |
|  | BACHD | Male | **3.007** |
|  |  | Females | 0.855 |
| REM | WT | Male | 1.446 |
|  |  | Females | 1.446 |
|  | BACHD | Male | **3.757** |
|  |  | Females | 1.736 |
| NREM | WT | Male | 0.860 |
|  |  | Females | 1.578 |
|  | BACHD | Male | **3.057** |
|  |  | Females | 1.950 |

**Fig 1.**
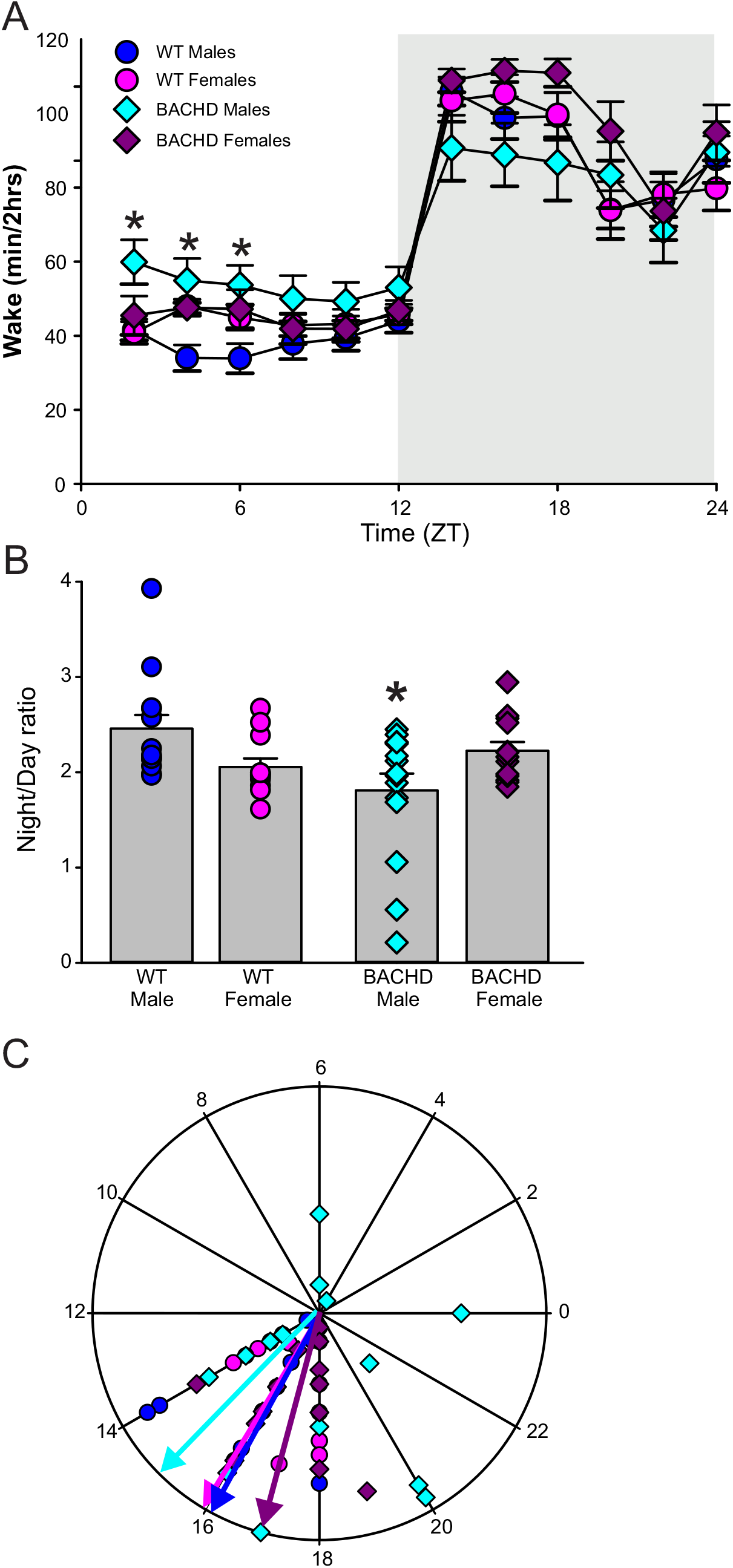
Temporal Pattern of Wake in WT and BACHD mice. EEG recordings were conducted in undisturbed mice in a 12:12 light/dark (LD) cycle over 24-hr. The data were analyzed using a three-way ANOVA with genotype, sex and time as factors. We found significant differences for genotype and time but not sex (**Table 2, 3**). (**A**) Waveform showing the daily rhythms of wake in the four groups. Significant differences as found by multiple comparison procedures (Holm-Sidak method) are indicated by asterisk. For this and the other figures, the gray-shaded area represents the dark (active) phase. Data are presented as mean ± SEM. (**B**) Amplitude of the daily rhythms in wake was estimated as the ratio of the minutes of wake during the night over the minutes of wake during the day. (**C**) The peak phase of the rhythm in wake was plotted on a polar display with the numbers on the axis indicating the time of day (ZT). The vector indicates the mean peak phase of each group. The male BACHD were significantly different in phase (**Table 4**) and exhibit higher variability than the other groups (**Table 5**).

Overall, male BACHD mice exhibited more wake during the day (resting period) and less during the night (active period) along with a high variability in phase compared to the other groups.

### REM sleep

Next, we carried out a similar analysis on the temporal patterns of REM sleep in the two genotypes. There were sex differences in the total amount of REM sleep (**Table 1**) with the females exhibiting more REM sleep than the males. This effect was more robust in the WT than in the BACHD mice. Next, we examined the rhythm in REM sleep using a three-way ANOVA with genotype, sex and time (day, night) as factors. We found significant differences for sex and time but not genotype (**Table 2**). Examination of the waveform of this cohort using 2-hr bins found that the REM levels of the two sexes were generally similar (**Fig. 2A, B**) although the 3-way ANOVA did detect significant differences in genotype, sex, and time (**Table 3**). We did not see any difference in the amplitude of the REM rhythms between the groups (**Fig. 2C, Table 4**). Finally, the groups exhibited a fair amount of variation in phase (**Fig. 2D**) with a significant interaction between genotype and sex (**Table 4**). Again, the BACHD males exhibited a great deal of variability in phase (**Table 5**) compared to the other 3 groups. Overall, sex differences in REM sleep were more evident in WT mice, while the male BACHD mice displayed more variability in phase compared to the other groups.

**Fig 2.**
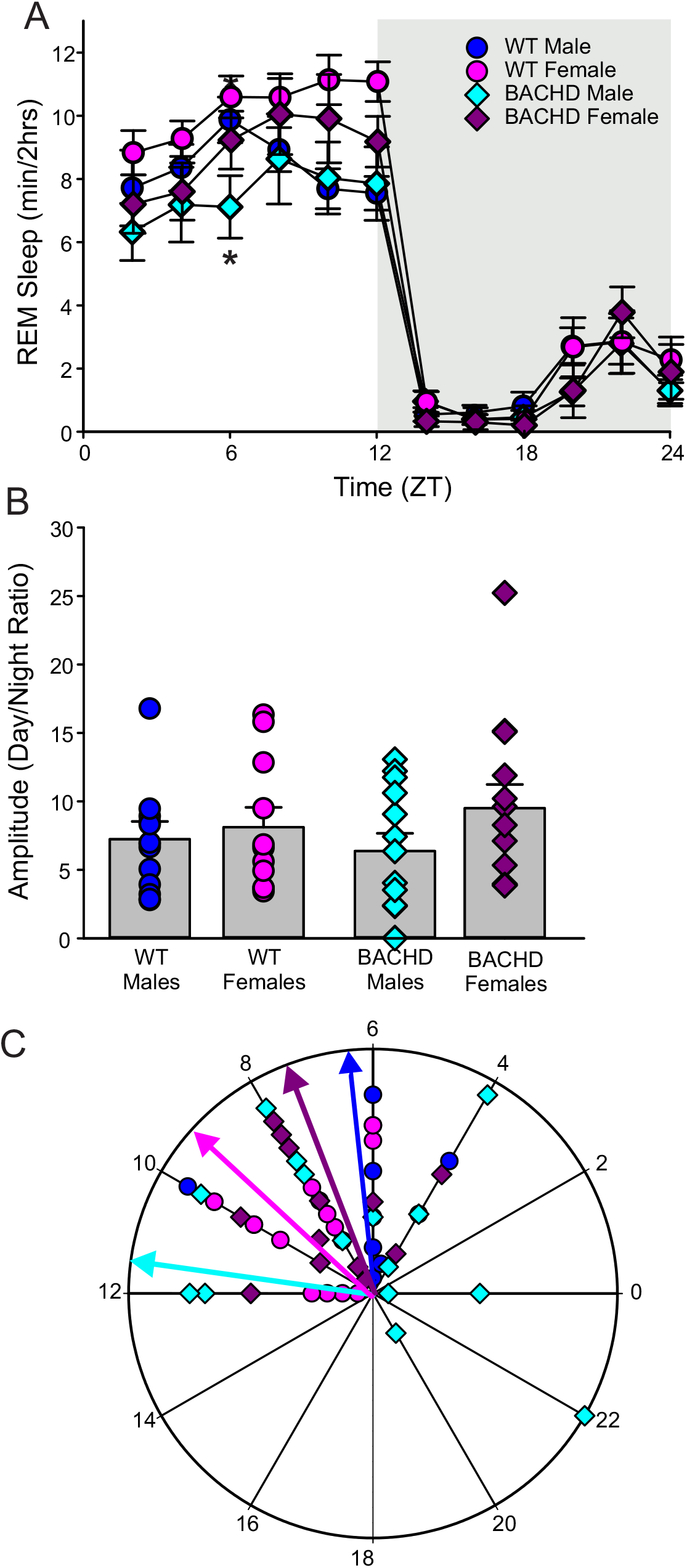
Temporal Pattern of REM in WT and BACHD mice. The data were analyzed using a three-way ANOVA with genotype, sex, and time as factors. While the daily patterns were broadly similar, there were significant differences in genotype, sex, and time (**Table 2, 3**). (**A**) Waveform showing the daily rhythms of REM sleep in the four groups. (**B**) Amplitude of the daily rhythms in REM was estimated as the ratio of the minutes of REM during the day over the minutes of REM during the day. (**C**) The peak phase of the rhythm in REM was plotted on a polar display with the numbers on the axis indicating the time of day (ZT). The vector indicates the mean peak phase of each group. There were no significant differences in phase (**Table 4**) but the male BACHD exhibited higher variability in peak phase than the other groups (**Table 5**).

### NREM sleep

Based on our prior data and the literature, we were particularly interested in examining potential sex differences in the temporal pattern of NREM sleep in the two genotypes. Sex had a significant effect on the total amount of NREM sleep (**Table 1**) with WT females exhibiting less NREM sleep than males. BACHD mice showed a similar trend, but the differences were not significant. The rhythm in NREM sleep was analyzed using a three-way ANOVA with genotype, sex, and time (day, night) as factors. Both sex and time, but not genotype, significantly influenced the rhythms (**Table 2**). Examination of the waveform of this cohort using 2-hr bins found that the NREM levels of the two genotypes were quite distinct in male mice with BACHD males exhibiting less sleep (**Fig. 3A)**. These sex differences were not seen between the females **(Fig. 3B**). The three-way ANOVA detected significant differences in all three variables (**Table 3**). The amplitude of the NREM rhythms was significantly reduced in males (**Fig. 3C, Table 4**). Furthermore, there was a significant interaction between genotype and sex detected in phase (**Fig. 3D, Table 4**) with the BACHD males exhibiting a great deal of variability in phase (**Table 5**). Finally, we carried out an additional analysis by assessing sleep fragmentation through measuring the number of NREM bouts and their duration (**Fig. 4**). The three-way ANOVA detected significant effects of genotype, sex, and time (**Table 6**). The male BACHD were most impacted and displayed a higher sleep fragmentation during the day with significant increases in the number of NREM bouts coupled with a decrease in their duration (**Fig. 4A, C**). Overall, sex differences in NREM sleep were more of a feature in WT mice. Male BACHD mice exhibited NREM sleep rhythms that were more fragmented and more variable in phase compared to the other groups.

**Table 6:** Analysis of NREM fragmentation using three way ANOVA with genotype (WT, BACHD), sex (male, female), and time (day, night) as factors. Interactions reported for genotype x sex x time.

| NREM | Genotype | Sex | Time (day, night) | Interactions |
| --- | --- | --- | --- | --- |
| Bout # | $F_{(1, 107)}=11.922, P < 0.001$ | $F_{(1, 107)}=9.947, P = 0.002$ | $F_{(1, 107)}=554.606, P < 0.001$ | $F_{(1, 107)}=4.924, P = 0.029$ |
| Bout Duration | $F_{(1, 107)}=7.101, P = 0.009$ | $F_{(1, 107)}=5.315, P = 0.023$ | $F_{(1, 107)}=0.069, P = 0.809$ | $F_{(1, 107)}=1.220, P = 0.272$ |

**Fig 3.**
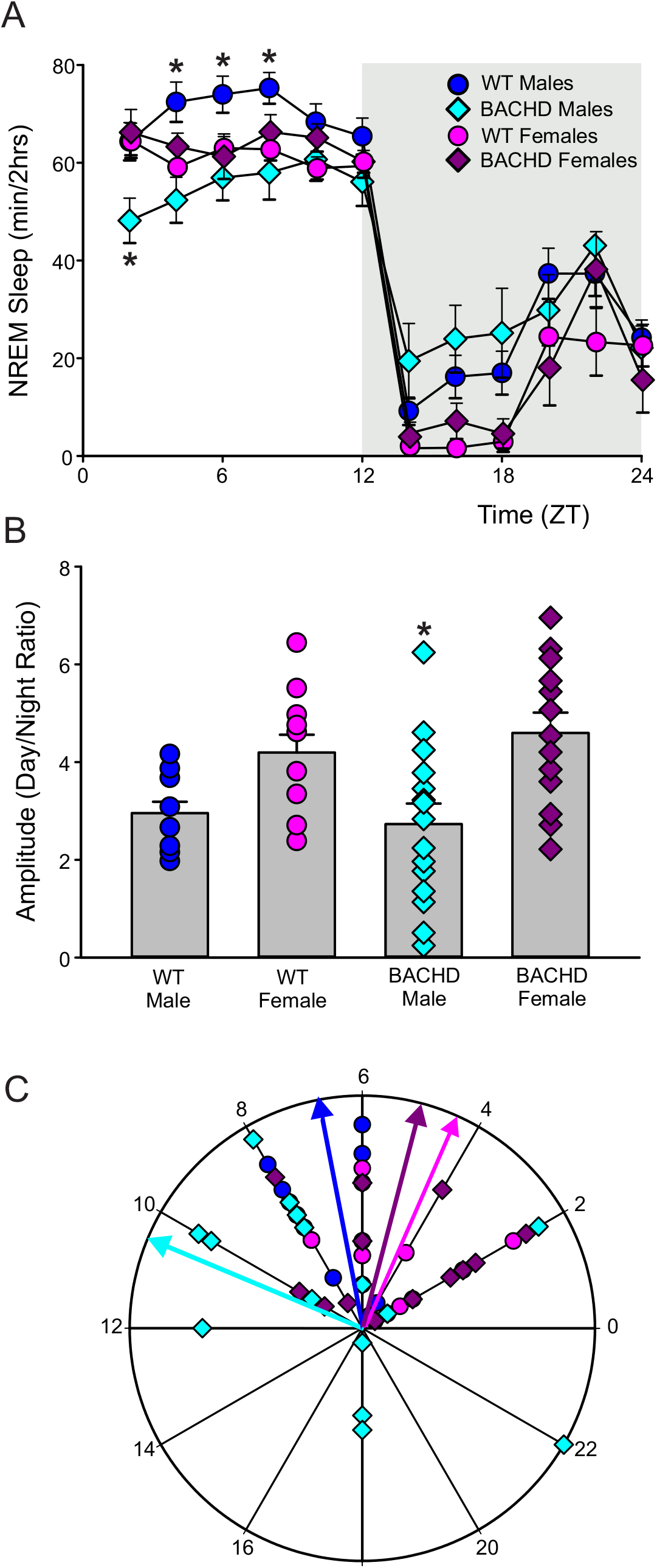
Temporal Pattern of NREM in WT and BACHD mice. The data were analyzed using a three-way ANOVA with genotype, sex, and time as factors. While the daily patterns were broadly similar, there were significant differences in genotype, sex, and time (**Table 2, 3**). (**A**) Waveform showing the daily rhythms of NREM sleep in the four groups. (**B**) Amplitude of the daily rhythms in NREM was estimated as the ratio of the minutes of NREM during the day over the minutes of NREM during the day. (**C**) The peak phase of the rhythm in NREM sleep was plotted on a polar display with the numbers on the axis indicating the time of day (ZT). The vector indicates the mean peak phase of each group. There were no significant differences in phase (**Table 4**) but the male BACHD exhibited higher variability in peak phase than the other groups (**Table 5**).

**Fig. 4.**
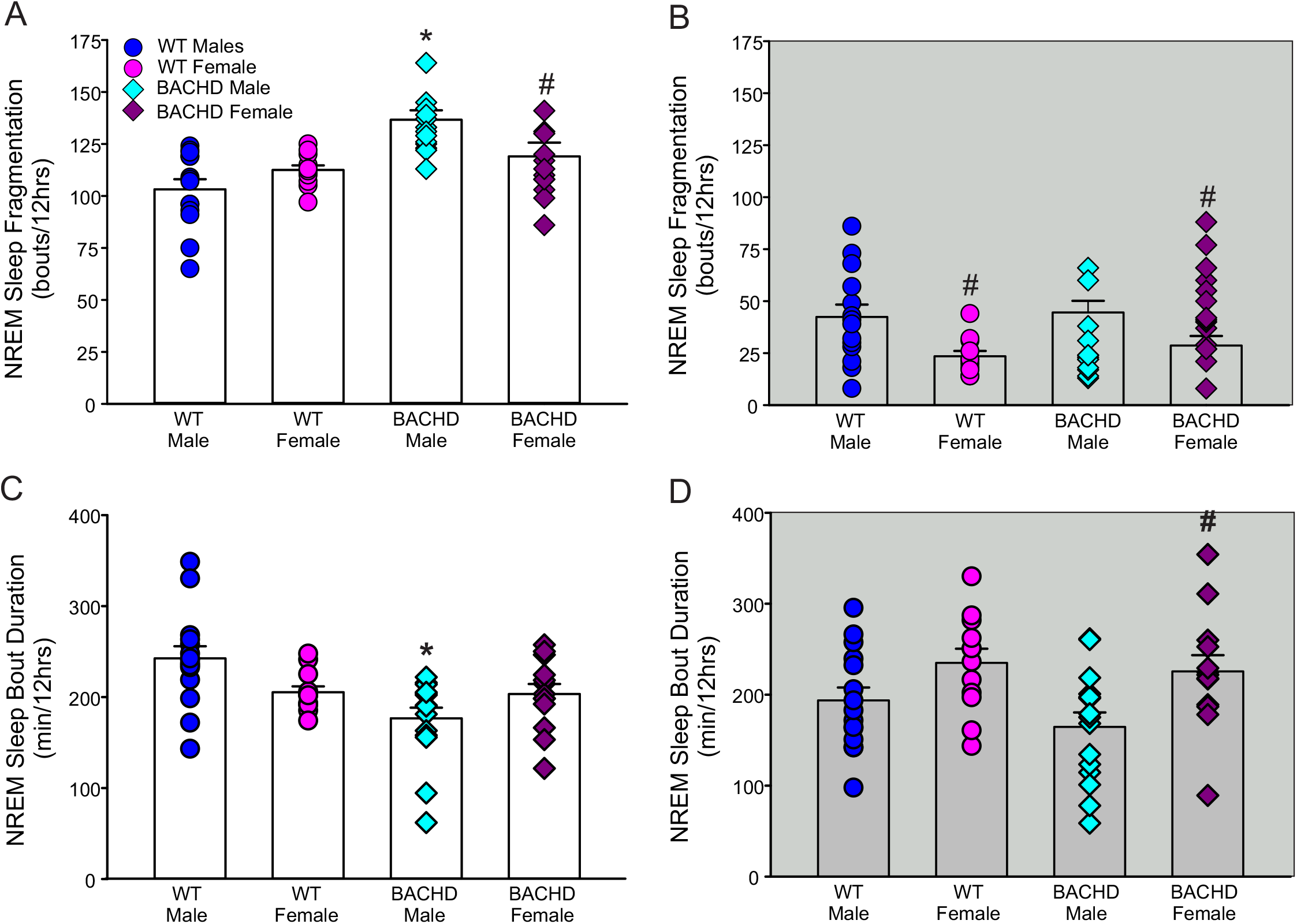
Sleep fragmentation increased in male BACHD mice. Measures of sleep fragmentation during NREM sleep were calculated from 24-hr EEG recordings. The three way ANOVA found significant differences for genotype, sex, and time (**Table 6**). Significant differences between sexes are indicated by an asterisk while a genotypic difference is shown with a cross-hatch. The number of NREM bouts were measured during the **(A)** day and **(B)** night. The average duration of NREM bouts during the **(C)** day and **(D)** night are also shown. In the day, male BACHD mice exhibited increased number of bouts that were also shorter in duration in the day. In the night, the female WT mice exhibited few bouts.

### EEG spectral power

It has been reported that HD mouse models display characteristic changes to the EEG spectra (Fisher *et al*., 2013; Kantor *et al*., 2013; Fisher *et al*., 2016), which recapitulate those displayed by HD patients (Leuchter *et al*., 2017). To investigate differences in the EEG spectra, we quantified the power values in the frontoparietal cortical region during NREM sleep. An analysis of the power spectral curves (1-40Hz) found evidence for sex differences in each genotype both during the day and night (**Table 7**). With this data, genotypic differences were predominant during the night (**Fig. 5**). During NREM sleep, the delta power (0.5-4 Hz) serves as an indicator of sleep pressure (Borbely and Tobler, 1989). We found evidence that male BACHD mice had reduced delta power compared to WT mice primarily during the night (**Fig. 5B**). Interestingly, female BACHD mice had more delta power during the night (**Fig. 5D**).

**Table 7:** analysis of power spectral curves with genotype, sex (male, female) and frequency (1-40Hz) as factors.

| Time | Genotype | Sex | Frequency (Hz) | Interactions |
| --- | --- | --- | --- | --- |
| Day | $F_{(1, 1679)}=1.462, P = 0.227$ | $F_{(1, 1679)}=32.860, P < 0.001$ | $F_{(39, 1679)}=87.306, P < 0.001$ | $F_{(39, 1679)}=1.269, P = 0.129$ |
| Night | $F_{(1, 1679)}=0.001, P=0.986$ | $F_{(1, 1679)}=41.793, P < 0.001$ | $F_{(39, 1679)}=95.320, P < 0.001$ | $F_{(39, 1679)}=1.220, P = 0.167$ |

**Fig. 5:**
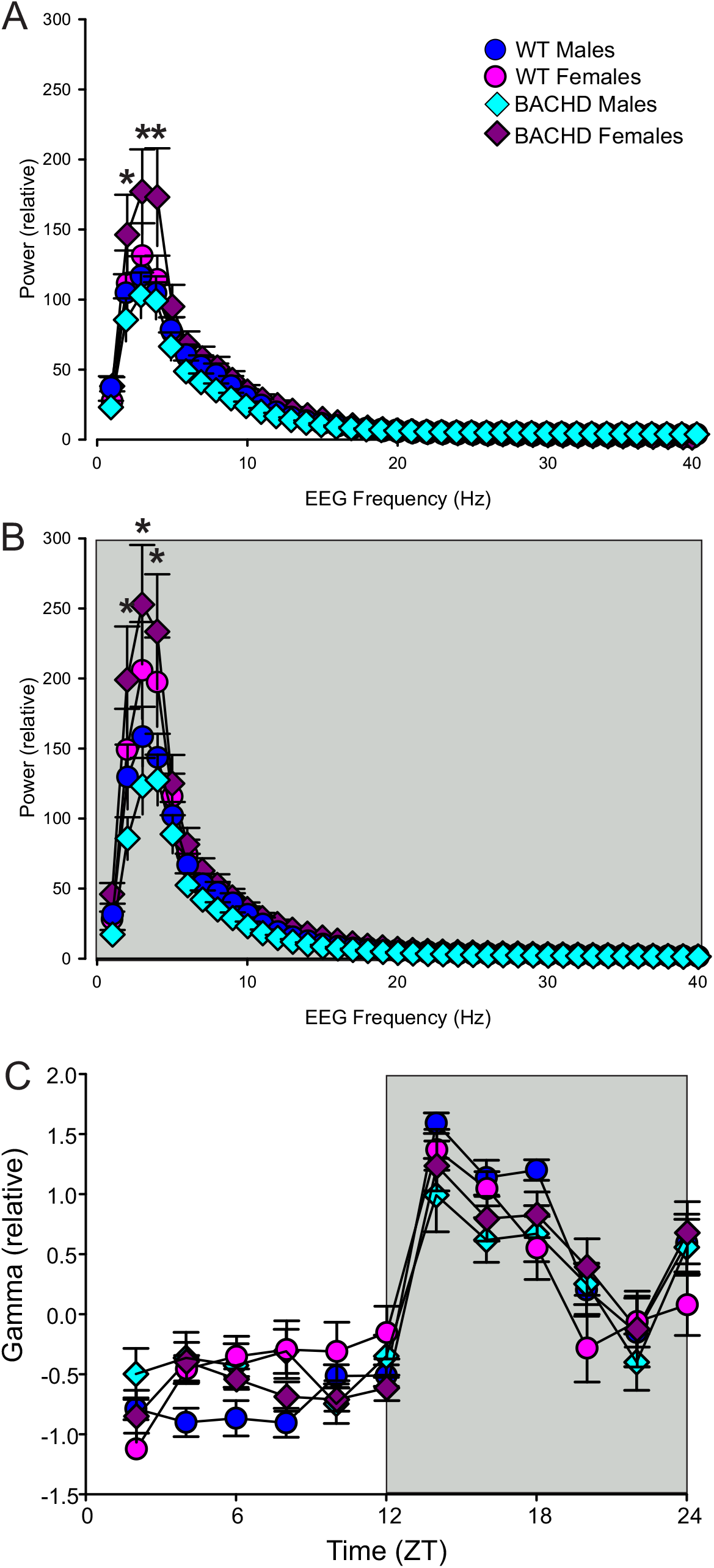
Altered EEG patterns in BACHD mice. Power spectral analysis was performed by applying a fast Fourier transform to raw 24h EEG waveforms. The data were analyzed using a three-way ANOVA with genotype, sex, and frequency as factors in the **(A)** day and **(B)** night. While the daily patterns were broadly similar, significant differences were detected for genotype, sex, and time (**Table 7**). For both sexes, the delta frequencies (1-4Hz) were most impacted by the mutation. We also examined the 24h pattern of gamma power across sleep/wake states in males and females (**C**).

Gamma power (20-40 Hz) did not clearly differ with sex or genotype in this data set (**Fig. 5, F**) although there were clear effects of time (3-way ANOVA: F_(11, 515)_ =41.389, P<0.001). The analysis of the spectral distribution of the EEG raises the possibility that there is an impairment in the sleep homeostatic system under undisturbed conditions in male, but not female, BACHD mice.

### Recovery from sleep deprivation

The most direct test of sleep homeostatic mechanisms is to examine the response to sleep deprivation. Therefore, we examined 18-hr of recovery sleep following 6-hr of sleep deprivation (SD) from ZT 0-6. First, the waveforms for each sleep state (**Fig. 6**) were analyzed using a 3-way ANOVA with genotype, sex, and time (2-hr bins) as factors. We found significant sex differences for REM and NREM sleep (**Table 8**). The SD protocol was equally successful in all groups and all groups exhibited a sleep rebound after the SD (**Fig. 6**). Analysis of recovery sleep between ZT 6-12, showed a strong increase in sleep (F_(1, 105)_=23.486, P < 0.001) but no difference between genotypes or sexes. While there was more variability in the response of male BACHD mice to SD; overall, the sleep homeostatic process appeared to be functional in both sexes of the mutant mice.

**Fig. 6.**
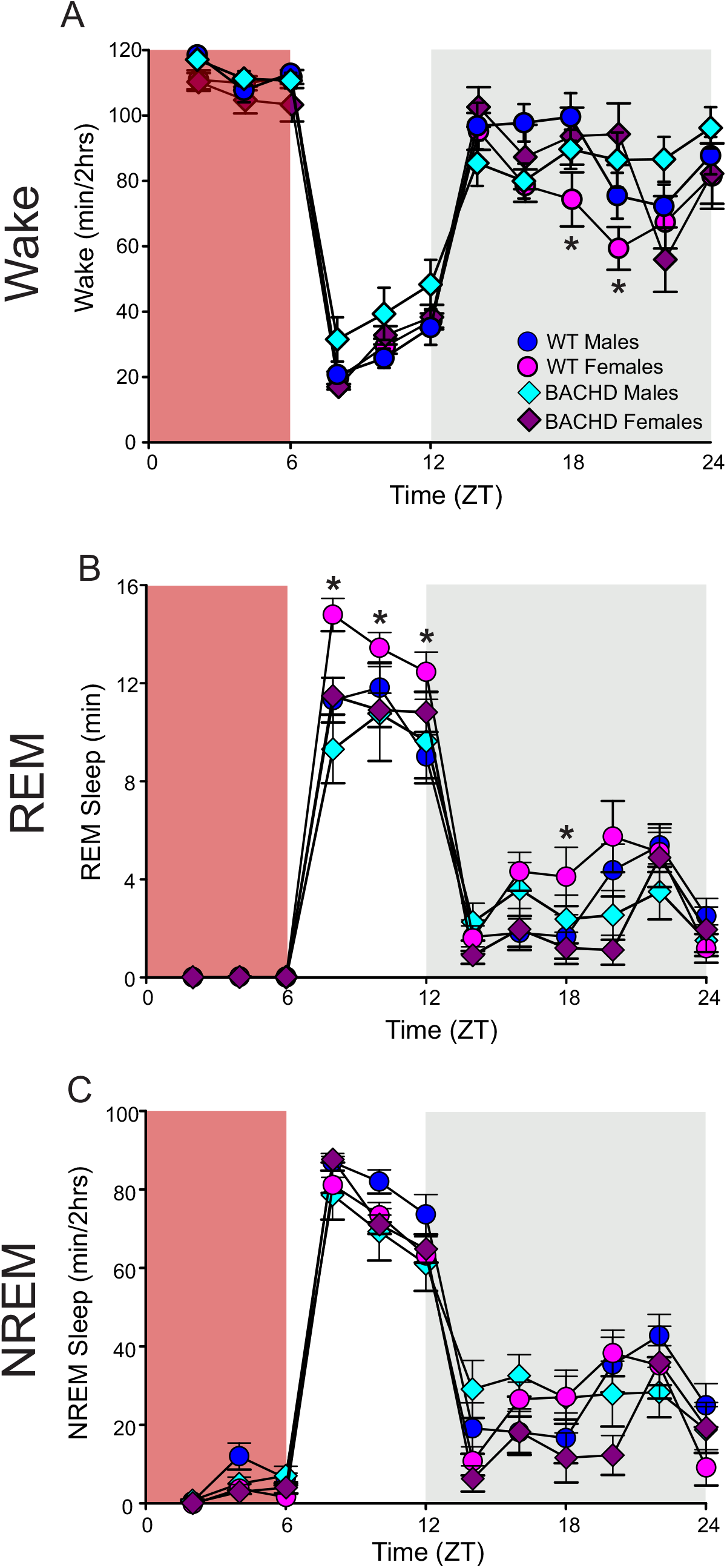
Sleep homeostatic mechanisms in BACHD mice. Mice were exposed to sleep deprivation (SD) at the beginning of their inactive phase (ZT0-6) using a gentle-handling protocol. The red-shaded area represents the time of SD. The amounts of **(A)** wake, **(B)** REM, and **(C)** NREM sleep in 2-hr bins are plotted. As analyzed by three-way ANOVA, all states exhibited differences in time, with REM and NREM sleep also exhibiting significant effects of genotype and sex (**Table 8**).

**Table 8:** Analysis of waveforms in response to sleep deprivation (SD) using three way ANOVA with genotype (WT, BACHD), sex (male, female), and time (2 hr bins) as factors. Interactions reported for genotype x sex x time.

| State | Genotype | Sex | Time (2 hr bins) | Interactions |
| --- | --- | --- | --- | --- |
| Wake | $F_{(1, 575)}=8.319, P = 0.004$ | $F_{(1, 575)}=0.927, P = 0.336$ | $F_{(11, 575)}=113.927, P < 0.001$ | $F_{(11, 575)}=2.828, P = 0.001$ |
| REM | $F_{(1, 625)}=14.680, P < 0.001$ | $F_{(1, 625)}=4.969, P = 0.026$ | $F_{(11, 625)}=123.097, P < 0.001$ | $F_{(11, 625)}=1.358, P = 0.189$ |
| NREM | $F_{(1, 625)}=3.134, P = 0.077$ | $F_{(1, 625)}=5.721, P = 0.017$ | $F_{(11, 625)}=141.404, P < 0.001$ | $F_{(11, 625)}=3.273, P < 0.001$ |

## DISCUSSION

HD patients often exhibit sleep and circadian disruptions before the onset of motor and cognitive symptoms (Fifel and Videnovic, 2020; Colwell, 2021). Similar disruption in the activity rhythms have been documented in a range of models including sheep (Morton et al., 2014), mini-pigs (Rieke et al., 2019), rodents (Morton et al. 2005; Bode et al. 2009; Kudo et al. 2011; Oakeshott et al. 2011; Loh et al. 2013; Woodard et al., 2017; Gu et al., 2022), and even *Drosophila* (Farago et al., 2019; Xu et al., 2019). One limitation of activity measures is that the deficits in the rhythms could be driven by motor dysfunction, thus it is important to also demonstrate the HD-driven deficits using physiological measures. In rodent models, EEG determinant of sleep help address this concern about activity measures. A number of sleep deficits have been documented, including increased wake during the inactive phase, increased sleep during the active phase, and loss of consolidated sleep (Fisher *et al*., 2013; Kantor *et al*., 2013; Loh *et al*., 2013; Fisher *et al*., 2016). In our present study, we found differences between the genotypes (BACHD vs. WT) in several key sleep parameters including time in wake (**Table 1**) and the daily waveform (2-hr bins) in wake, REM, and NREM states (**Fig. 1-3**). Sleep fragmentation was also evident with increases in the total number of NREM bouts as well as a reduction in the bout duration (**Fig. 4**). Our findings demonstrated clear dysfunction in the sleep/wake architecture and sleep fragmentation in young adult BACHD mice compared to WT controls.

EEG signal is generated by cortical brain activity, and rhythms in EEG oscillations are produced through interactions between thalamic and cortical neurons (Feyissa and Tatum, 2019). Neural oscillations occur in different frequency bands, (delta, 0.5-4 Hz; theta, 4-8 Hz: alpha, 8-14 Hz; beta, 14-20 Hz; gamma, >20 Hz), each with distinct functional characteristics (Saby and Marshall, 2012). Aberrant EEG patterns have been observed in many different nervous system disorders (e.g. Han et al., 2013; Fitzgerald and Watson, 2018; Horvath et al., 2018; Sebastián-Romagosa et al., 2020). Specifically, in HD patients, there is evidence for abnormal EEG power associated with specific frequency bands, with changes in alpha, beta and delta commonly noted (Wiegand et al., 1991; Bylsma et al., 1994; Hunter et al., 2010; Painold et al., 2010; Piano et al., 2017). Similarly, in transgenic HD sheep, EEG recordings found a significant decrease in delta (0.5-4 Hz) power across the night compared to normal sheep (Vas et al., 2021). Finally, several studies have found characteristic changes in the EEG spectra in mouse models including the R6/2 (Fisher et al., 2013; Kantor et al., 2013), the R6/1 (Lebreton et al., 2015), and Q175 (Fisher et al., 2016). Based on this prior data, we expected to see alterations to the EEG spectra, but genotypic differences did not emerge out of our data (**Table 7**). Based on the data from other studies, it seems likely that differences in EEG spectra would emerge as the symptoms progress in the BACHD model.

In prior work, we have shown that male BACHD mice exhibit more severe deficits in activity rhythms and motor coordination than females (Kuljis *et al*., 2016). The focus of the present study was to explore possible sex differences in the sleep/wake cycle and EEG power spectrum of BACHD mice. We found sex differences in the total amount of REM and NREM sleep (**Table 1**) but also in the temporal pattern of these states (**Fig. 2, 3, Table 3**). Males BACHD mice exhibited reduced amounts of NREM in the day, a feature which is not seen in the female BACHD (**Fig. 3**). Additionally, the male BACHD mice exhibited a striking increase in variability in the phase of the rhythms in each state (**Table 5**). Overall, the EEG measures of state are consistent with our prior behavioral data indicating the male BACHD mice are more impacted than female BACHD mice.

NREM delta power serves as a biomarker of homeostatic sleep drive, as delta power is enhanced after prolonged wakefulness, but declines as sleep deepens (Borbely et al., 1981; Long et al., 2021). Based on the reduced sleep in the mutants, we expected to see higher delta levels during NREM. Surprisingly, female, but not male, BACHD conformed to these expectations (**Fig. 5**). Prior work has also found reduced delta wave power in male mouse models including the R6/2 (Fisher et al., 2013; Kantor et al., 2013), the R6/1 (Lebreton et al., 2015), and Q175 (Fisher et al., 2016). To further test sleep homeostatic mechanisms, we examined how the 12 week-old BACHD and WT mice responded to a 6-hr sleep deprivation protocol. All groups were effectively sleep deprived and all groups responded to SD with an increase in sleep amount (**Fig. 6**). In the R6/2 and Q175 models (Fisher et al., 2013; 2016), the responses to SD were largely intact although some genotypic differences were seen in the R6/2 mice at the end of the disease progression. Our analysis did uncover significant genotype and sex differences in the homeostatic response to SD (**Table 8**).

HD is a genetic disorder with the mutation on an autosomal chromosome, so ultimately comparable numbers of males and females will develop HD. However, this does not mean that the trajectories of the disease are the same (Zielonka and Stawinska-Witoszynska, 2020) and interrogation of human datasets suggests that females are more vulnerable (e.g. Hentosh et al., 2021). In preclinical models, the data from the CAG 140 (Dorner et al., 2007), Q175 (Padovan-Neto et al., 2019), and the BACHD mouse model (Kuljis et al., 2016) all suggest that males being more impacted early in the disease trajectory. Mechanistically, the sex differences are likely to be driven by hormonal factors. In preclinical models, 17β-estradiol protects against β-amyloid toxicity (Guerra *et al*., 2004), and plasma 17β-estradiol was found to be correlated with striatal neuron loss in male HD rats (Bode *et al*., 2008). Our findings suggest that females are protected from sleep/wake deficits early in the HD progression. Based on this study’s findings, we believe that an examination of sex differences in prodromal non-motor phenotypes is warranted. It is worth point out that new treatment strategies may emerge from exploring the impact of estrogen/ERα signaling in HD (Yilmaz et al., 2019; Veenman, 2020; Koszegi and Cheong, 2022; Tecalco-Cruz et al., 2023).

## Acknowledgments

We thank the team who helped us with the sleep deprivation experiments and animal maintenance including Dante Dikeman, Mary Dover, Kyle Golden, Cameron Jackson, Noah Liberty, Melika Madani, Giselle Melendez, Damion Trotter, Alexis Tucker, Scott Vincent, and Sam Wimmer. In addition, we would thank the CHDI Foundation for providing the backcrossed BACHD mice used to establish the breeding colony. Finally, we would also thank Dr. K. Tamai for her work managing the breeding colony and study area used for these experiments.

## Author Contributions

All authors had full access to all the data in the study and take responsibility for the integrity of the data and the accuracy of the data analysis. Conceptualization, K.N.P., C.A.G., C.S.C.; Methodology, E.C., K.N.P., Investigation, E.C.; K.Z.; G.S.; Formal Analysis, E.C.; Writing – Original Draft, E.C.; Writing – Review & Editing, K.N.P, C.A.G., C.S.C.; Visualization, E.C., C.S.C; Supervision, K.N.P, C.A.G, C.S.C.; Funding Acquisition, K.N.P., C.S.C.

The authors have no conflict of interest to declare.

